# RuHere (Are You Here?): An R package to obtain, validate, and clean species records using metadata and specialist range information

**DOI:** 10.64898/2026.02.02.703373

**Authors:** Weverton C. F. Trindade, Fernanda S. Caron

## Abstract

1. Species occurrence data are fundamental to understanding, predicting, and conserving global biodiversity. However, biodiversity datasets remain affected by substantial data-quality issues, particularly erroneous or imprecise geographic coordinates. Most available tools for identifying problematic records rely primarily on automated spatial or metadata-based checks and rarely integrate expert-curated species range information, which can reveal introductions or geographic errors that often escape standard validation procedures.
2. Here, we introduce RuHere, an R package designed to manage species occurrence data, flag potential errors, and support the iterative exploration of problematic records. RuHere streamlines the data-cleaning process by integrating six main steps: (1) obtaining species occurrence records; (2) merging datasets and standardizing spatial information; (3) flagging records based on metadata; (4) flagging records using expert-derived distribution data; (5) visualizing, investigating, and summarizing flagged issues in the final datasets; and (6) exploring and reducing sampling bias.
3. We demonstrate the applicability of RuHere using occurrence data for a plant species (*Araucaria angustifolia*) and an animal species (*Cyanocorax caeruleus*). Nearly 75% of records were flagged as potentially problematic, including records identified exclusively by functions relying on specialist range information.
4. The main strengths of RuHere lie in its integrated and computationally efficient workflow, its tools for exploring and evaluating flagged records, and its ability to incorporate expert-derived distribution data to identify occurrences outside a species’ known natural range. By combining metadata-based checks, coordinate validation, and specialist knowledge, RuHere provides a robust and reproducible framework for improving the quality of species occurrence datasets.

## 1. INTRODUCTION

Species occurrence data are fundamental to understanding, predicting and conserving global biodiversity (Peterson et al., 2018). Over the past two decades, the availability of such data has increased exponentially (Telenius, 2011), with the Global Biodiversity Information Facility (GBIF) now providing more than 3.5 billion occurrence records (as of January 2026). Despite this, biodiversity datasets remain affected by substantial data-quality issues, including erroneous or imprecise geographic coordinates (Zizka et al., 2020). For instance, Trindade & Marques (2023) found that 74% of occurrence records of native plants from the Atlantic Forest Hotspot are problematic and unsuitable for biodiversity analyses. Thus, improving the accuracy and reliability of primary biodiversity data is essential for understanding our planet’s biodiversity and addressing global environmental issues.

Data cleaning, the process of identifying and removing problematic records, has become a mandatory step in biodiversity research (Sillero et al., 2021). While manual cleaning may be feasible for small, well-known datasets, it is impractical for large-scale analyses (Zizka et al., 2019). Indeed, data preparation is usually the most time-consuming process in studies relying on occurrence data (Sillero et al., 2021). Subjective manual decisions can further amplify existing biases, potentially distorting biodiversity patterns. To address these limitations, several R packages automate data cleaning, such as CoordinateCleaner (Zizka et al., 2019), plantR (de Lima et al., 2021), and bdc (Ribeiro et al., 2022). However, a critical gap remains: most tools do not integrate expert-curated species range information, which can reveal introductions or geographic errors that often escape automated spatial tests (Reginato & Michelangeli, 2020).

Here, we present RuHere, an R package designed to manage species occurrence data, flag potential errors, and support the iterative exploration of problematic records. RuHere streamlines the data-cleaning workflow by combining automated spatial filters, metadata validation, and direct integration with specialist range information. The package provides a comprehensive, transparent, and reproducible framework, from the initial acquisition and merging of data from multiple repositories (GBIF, SpeciesLink, BIEN, and iDigBio) to the mitigation of sampling bias in geographic and environmental space. By offering researchers greater flexibility and control over the data-cleaning process, RuHere ensures that biodiversity data are reproducible, high-quality, and fit for use in ecological and conservation applications.

## 2. DESCRIPTION

### 2.1 Overview

The RuHere package provides a comprehensive workflow designed to streamline the acquisition, standardization, and validation of species occurrence data. The proposed workflow is structured into six integrated steps (Figure 1): (1) obtaining species occurrence records; (2) merging datasets and standardizing spatial information; (3) flagging records using metadata (i.e., associated information); (4) flagging records using specialists’ range information; (5) exploring and reducing sampling bias; and (6) visualizing, investigating, and summarizing flagged issues within the final datasets. To preserve data integrity, RuHere adopts a “flag-first, remove-later” approach, in which each validation function appends a logical column to the dataset indicating whether a record passed (TRUE) or failed (FALSE) a specific test. This strategy ensures full reproducibility and user control, resulting in a refined, fit-for-use dataset accompanied by comprehensive reports and diagnostic figures that document the data-cleaning process. Further details on each step of the workflow are provided below. In addition, extensive documentation and tutorials for all functions are available on the package website (https://wevertonbio.github.io/RuHere).

**Figure 1.**
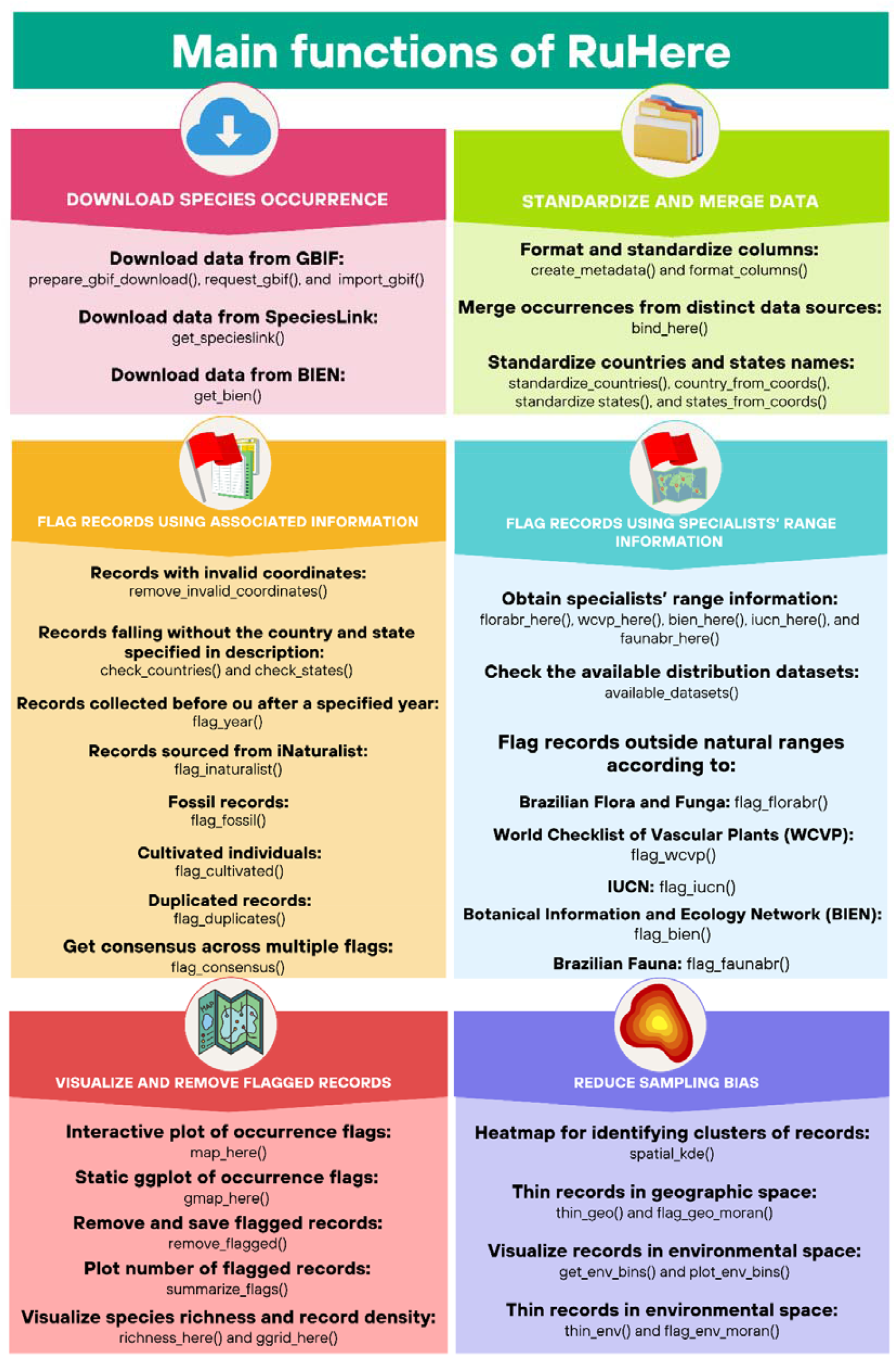
Main functions of the workflow proposed in RuHere for obtaining, validating, and cleaning species occurrence records using metadata and specialists’ range information.

### 2.2. Downloading species occurrence

Integrating data from multiple biodiversity repositories poses two major challenges: data acquisition and format standardization. Each database adopts distinct data structures and column naming conventions, which complicates the direct merging of datasets. RuHere addresses these challenges by enabling users to download and harmonize species occurrence records directly within the R environment, performing an initial standardization of key fields at the time of data acquisition. The package currently accesses data from four major repositories: GBIF (https://www.gbif.org/), SpeciesLink (https://specieslink.net/), BIEN (Botanical Information and Ecology Network; https://bien.nceas.ucsb.edu/bien/), and iDigBio (Integrated Digitized Biocollections; https://www.idigbio.org/). For services requiring authentication, such as GBIF and SpeciesLink, dedicated functions store API credentials securely within the user’s R environment, facilitating automated and reproducible download requests.

### 2.3. Standardizing and merging data

Following data acquisition, RuHere performs a standardized harmonization of occurrence records to ensure consistency across datasets originating from different repositories, adopting the Darwin Core (DwC) standard as a common data model (Wieczorek et al., 2012). Standardization is implemented through the *format_columns()* function, which harmonizes column names and data classes and corrects text encodings for fields that commonly contain special characters. This process relies on internal metadata templates provided for datasets from GBIF, SpeciesLink, BIEN, and iDigBio. RuHere also supports the integration of datasets from additional sources, such as data papers, by allowing users to define custom metadata templates using the *create_metadata()* function. These templates map original column names to the package’s standardized schema. Once standardized, occurrence records from multiple sources can be combined into a single dataset using the *bind_here()* function.

After initial harmonization and merging, species occurrence data often retain spatial inconsistencies that can compromise subsequent analyses. Common issues include different spellings or abbreviations of the same country or state/province (e.g., Brasil and Brazil), as well as missing administrative information. RuHere addresses these issues by standardizing country, state, and province names using the *standardize_countries()* and *standardize_states()* functions. These functions compare values in the original dataset (i.e., country and state/province fields) against a comprehensive dictionary of official names and postal codes. When country or state/province information is missing, the functions can optionally infer and fill these attributes based on the administrative units in which the geographic coordinates fall.

### 2.4. Flagging records using metadata

In addition to geographic coordinates, species occurrence datasets often contain valuable contextual information describing the conditions under which records were collected (Zizka et al., 2020), collectively referred to as metadata. RuHere provides a suite of tools that leverage metadata to identify potentially problematic records. Records with missing or invalid geographic coordinates can be identified and flagged using the *remove_invalid_coordinates()* function. After country and state names have been standardized, the *check_countries()* and *check_states()* functions can be used to evaluate whether each record’s coordinates fall within the country and state or province specified in the associated metadata.

Occurrence records may be taxonomically and spatially correct but still fail to represent the realized or contemporary niche of a species. This is particularly relevant in analyses that assume equilibrium between species distributions and environmental conditions (Sillero et al., 2021). RuHere addresses this issue through metadata-based flagging tools. The *flag_year()* function identifies records outside a user-defined temporal range, which is useful when modeling species distributions under contemporary climate conditions. Fossil records, which reflect past climatic and ecological contexts, are detected by the *flag_fossil()* function. For plant species, records of cultivated individuals that may occur outside the natural range or persist only under artificial conditions are flagged using the *flag_cultivated()* function. Some occurrence records, particularly those accessed through GBIF, originate from the citizen science platform iNaturalist (https://www.inaturalist.org/). Because many of these records are contributed by non-specialists, users may choose to treat them differently or exclude them from certain analyses. The *flag_inaturalist()* function allows users to flag all iNaturalist-derived records or only those not classified as Research Grade, offering great flexibility. Duplicate records can be identified using the *flag_duplicates()* function. RuHere allows users to define rules for retaining records among duplicates, for example by prioritizing the most recent occurrence or a specific data repository. Additionally, duplicates can be flagged at user-defined spatial resolutions by identifying records occurring within the same raster cell.

### 2.5. Flagging records using specialists’ range information

When a species is known to occur only within specific geographic regions, records falling outside these areas can be flagged as potential errors, misidentifications, or introductions (Reginato & Michelangeli, 2020). RuHere supports the cross-referencing of occurrence records against five authoritative datasets containing expert-curated distribution information. The World Checklist of Vascular Plants (WCVP) and the International Union for Conservation of Nature (IUCN) provide distribution information using botanical countries, corresponding to Level 3 of the World Geographical Scheme for Recording Plant Distributions, a reference map that is also included in the package. The BIEN provides range polygons for plant species derived from ecological niche models (Maitner et al., 2018). The Brazilian Flora and Funga (florabr) and the Taxonomic Catalog of the Brazilian Fauna (faunabr) are complementary initiatives that document the distribution of plant, fungal, and animal species occurring in Brazil (Trindade, 2024, 2025). Because these resources are updated regularly, users must download the most recent versions using the functions *wcvp_here(), iucn_here(), bien_here(), florabr_here()*, and *faunabr_here()*.

Once the datasets have been downloaded, RuHere provides the functions *flag_wcvp(), flag_iucn(), flag_bien(), flag_florabr()*, and *flag_faunabr()* to evaluate whether occurrence records fall within the known geographic range of each species. All functions allow the application of a user-defined buffer distance around distributional boundaries to account for spatial uncertainty. The *flag_consensus()* function allows users to derive a consensus across multiple flags. Consensus can be computed in two ways: a record may be considered valid only if all specified flags indicate validity, or if at least one specified flag indicates validity. For example, users can combine the results from *flag_bien()* and *flag_iucn()* to flag as invalid only those records that fall outside the natural ranges defined by both BIEN and IUCN.

### 2.6. Visualizing and removing flagged records

One of the main strengths of RuHere lies in its tools for exploring and evaluating flagged records. After records have been flagged, users can visualize their spatial distribution to assess whether identified issues are consistent with expected ecological and geographic patterns. RuHere provides two flexible functions for visualizing occurrence records. The *map_here()* function generates interactive maps, allowing users to dynamically explore flagged records (Figure 2c). In contrast, the *ggmap_here*() function produces static maps, which are particularly useful for summarizing results and reporting flagged patterns in figures (Figure 2a and 2b). When using *ggmap_here()*, users may either display all flagged records in a single map or generate multi-panel figures in which each panel corresponds to a different flag type.

**Figure 2.**
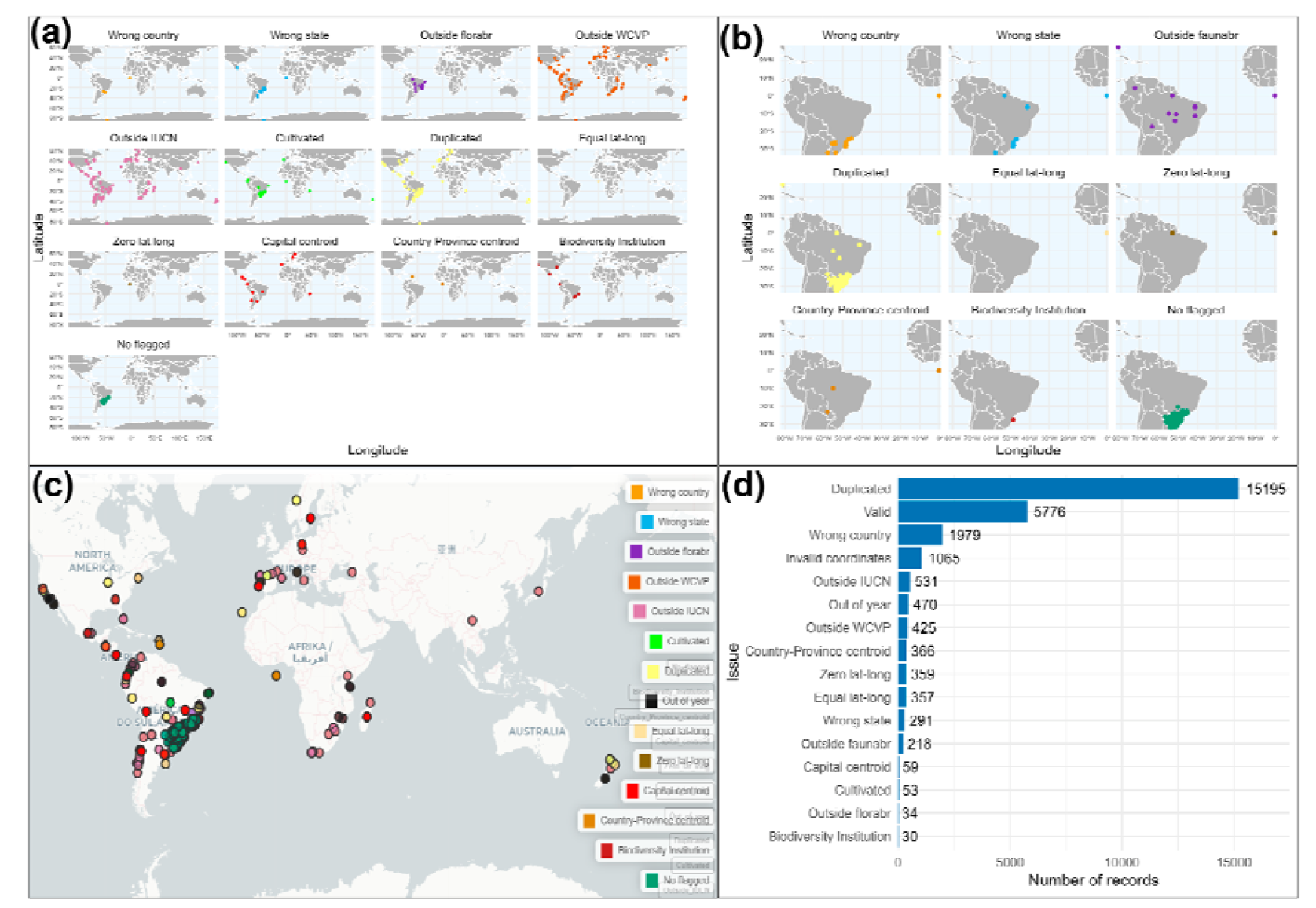
Outputs from RuHere flagging functions: (a) Paraná pine and (b) azure jay flagged records generated with *ggmap_here()*; (c) interactive map from *map_here()* for Paraná pine; and (d) bar plot from *summarize_flags()* summarizing the number of flagged records for both species.

Because RuHere adopts a “flag-first, remove-later” approach, the *remove_flagged()* function can be used to exclude records flagged as invalid. By default, the function considers all flags present in the dataset, but users may specify a subset of flags to apply. When a target folder is provided, the function automatically saves the removed records, labeled by the corresponding flag names. In addition, users can explicitly specify which flagged records should be retained or identify unflagged records to be removed. This flexibility allows researchers to incorporate their expert knowledge to complement automated data-quality checks.

Reporting the number of flagged and removed records is essential to ensure transparency and reproducibility. The *summarize_flags()* function can be used to generate a bar plot summarizing the number of records flagged by each flagging procedure (Figure 2d). In addition, the *richness_here()* and *ggrid_here()* functions allow visualization of the spatial distribution of species richness or record density across the study area. These visualizations are particularly useful for assessing how data-cleaning steps influence spatial patterns in the dataset.

### 2.7. Compatibility with CoordinateCleaner

CoordinateCleaner is an R package that performs automated tests to identify suspicious geographic coordinates (Zizka et al., 2019). Among its functions, it compares occurrence coordinates against reference databases of country and province centroids, country capitals, urban areas, and biodiversity institutions. RuHere is fully compatible with the flags generated by CoordinateCleaner, as both packages append logical flag columns to the dataset. Consequently, all RuHere functions for visualizing and summarizing flags can also be applied to the flags produced by the *clean_coordinates()* function in CoordinateCleaner.

### 2.8. Reducing sampling bias

Sampling bias is a common issue in primary biodiversity data, often producing spatial clusters that reflect accessibility and collector preferences rather than species ecology. In RuHere, users can explore such patterns using the *spatial_kde()* function, which generates occurrence density heatmaps based on kernel density estimation. A widely used strategy to reduce sampling bias is the thinning of occurrence records, which involves removing records that are spatially or environmentally close to each other (Aiello-Lammens et al., 2015). Thinning can be performed in geographic or environmental space, as both approaches improve the performance of species distribution models (Boria et al., 2014; Castellanos et al., 2019). RuHere supports both thinning strategies. In geographic space, *thin_geo()* retains a single record per species within d kilometers, allowing user-defined priority rules (e.g., most recent records). In environmental space, *thin_env()* divides environmental variables into n bins and flags records that fall within the same multidimensional environmental block as redundant. Retained records are flagged as TRUE in the *thin_geo* or *thin_env()* columns and can be filtered using *remove_flagged()*.

A central challenge in record thinning is determining an appropriate thinning distance or number of bins, since we rarely have sufficient biological justification for choosing a specific value (Lamboley & Fourcade, 2024). To address this issue, the *flag_geo_moran()* and *flag_env_moran()* functions build on the approach proposed by Velazco et al. (2021), computing spatial autocorrelation (Moran’s I) across datasets generated using a range of thinning distances or bin numbers and selecting the value that minimizes average spatial autocorrelation. We extend this procedure by introducing additional selection criteria that prevent the choice of datasets with too few records or unrealistically low Moran’s I values. To enhance performance, the core routine responsible for computing Moran’s I, *moranfast()*, was implemented in C++, substantially reducing computational time and memory usage.

A potential issue when filtering records in geographic space is that two nearby records may actually occur in distinct environmental conditions, especially in highly heterogeneous regions. Thinning in environmental space can suffer from the opposite issue of geographic thinning: environmentally similar records may be geographically far apart, potentially removing important information about the species’ geographic range. In such cases, we recommend using the *flag_consensus()* function to identify records that are redundant in both geographic and environmental space.

### 2.9. Empirical example: the Paraná pine and the azure jay

We demonstrated the RuHere applicability in obtaining and preparing occurrence data of the two species represented in the logo of the package: the Paraná pine (*Araucaria angustifolia*) and the azure jay (*Cyanocorax caeruleus*). The Paraná pine is an emblematic coniferous tree native to the subtropical highlands of Brazil, Argentina, and Paraguay, while the corvide azure jay is one of its main seed dispersers. Occurrence data for both species were retrieved from the databases supported by RuHere (GBIF, SpeciesLink, BIEN, and iDigBio) and complemented with records from an external source, the atlanticr R package (Vancine et al., 2025). We then identified potentially problematic records using flagging functions implemented in RuHere, as well as those provided by the CoordinateCleaner package. After removing flagged records, we mitigated sampling bias by combining thinning procedures in both geographic and environmental space using *flag_geo_moran()* and *flag_env_moran()*. The R scripts used in the analyses are available in Appendix S1.

## 3. RESULTS

We obtained a total of 22,885 occurrence records, comprising 6,204 records for the Paraná pine and 16,681 records for the azure jay. After identifying and removing problematic records, 5,776 occurrences were retained as suitable for biodiversity analyses, corresponding to an overall reduction of nearly 75%. The largest reduction occurred in the azure jay dataset, for which only 3,363 records (approximately 20% of the original data) passed all validation steps. In contrast, 2,413 records for the Paraná pine (39% of the total) were retained. The most frequent issues detected were duplicate records, occurrences with coordinates falling outside the country indicated in the metadata, and records with invalid geographic coordinates (Figure 2). Of the 17,109 flagged records, 458 were identified exclusively by functions that rely on specialist-curated distribution information from IUCN, WCVP, florabr, and the faunabr.

Most retained records were strongly concentrated around the city of Curitiba, the capital of Paraná State, particularly for the Paraná pine (Figure 3). During geographic thinning, a distance of 15 km was selected as the optimal threshold for reducing spatial autocorrelation while retaining a sufficient number of records. In environmental space, the optimal configuration corresponded to 80 bins for both species. After combining the two approaches and removing only records that were redundant in both geographic and environmental space, we retained 756 records for the Paraná pine and 942 records for the azure jay.

**Figure 3.**
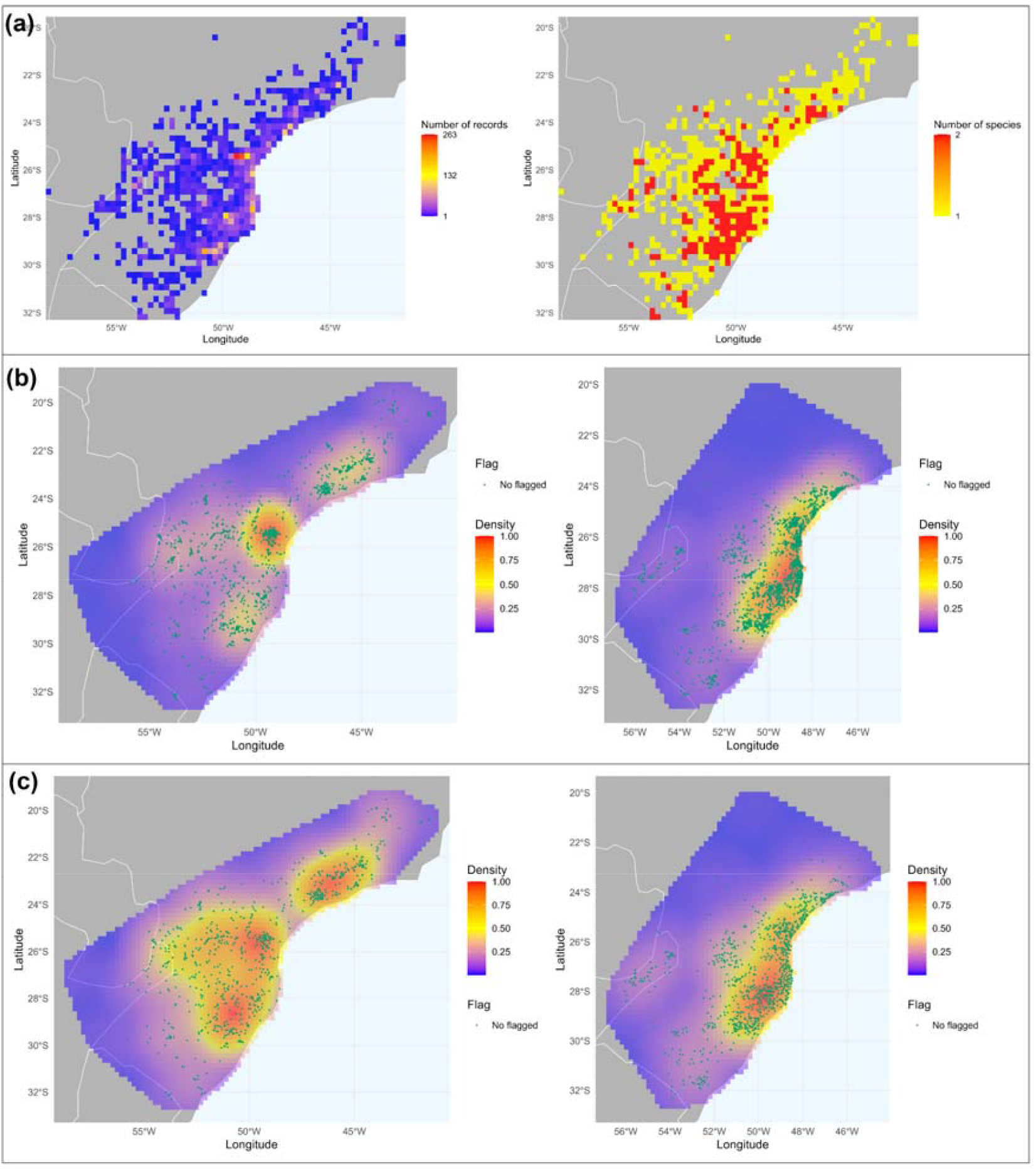
(a) Output of *ggrid_here()* showing the number of records and species in the raw dataset; (b) heatmaps of occurrence records for Paraná pine (left) and azure jay (right) before thinning; and (c) after thinning in geographic and environmental space.

## 4. DISCUSSION

Several R packages provide tools for cleaning biodiversity data. Some functionalities implemented in RuHere overlap with those available in existing packages, such as the identification of inconsistencies between occurrence coordinates and country information in the metadata, as implemented in bdc (Ribeiro et al., 2022); the detection of cultivated plant individuals in plantR (de Lima et al., 2021); the identification of duplicate records in CoordinateCleaner (Zizka et al., 2019); and geographic thinning of occurrence records in spThin (Aiello-Lammens et al., 2015). Despite these similarities, RuHere performs these tasks using a unified workflow built on computationally efficient tools, including data.table and terra R packages, and core routines rewritten in C++, resulting in improved speed and memory efficiency.

The main novelty of RuHere lies in its ability to incorporate specialist-curated range information to identify occurrence records falling outside a species’ known natural distribution, one of the most robust approaches to validating biodiversity data. In our case study, several records that were clearly outside the species’natural range were detected exclusively by functions relying on specialist range information. However, specialist-curated distribution data remain scarce for many taxonomic groups. For example, although IUCN provides detailed range polygons through its website, these data cannot currently be accessed programmatically via an R package and are therefore not directly integrated into RuHere. A planned improvement for future versions of the package is the inclusion of additional regional and taxon-specific databases that provide expert-derived distribution information, similar to the role played by florabr and faunabr for Brazilian taxa. We encourage user feedback and contributions to help address these limitations and guide the development of new features.

By integrating metadata-based checks, coordinate validation, and expert-derived distribution information within a single workflow, RuHere provides a transparent and reproducible framework for identifying potential issues in species occurrence datasets. In addition, the interactive visualization of flagged records supports the incorporation of expert knowledge alongside automated data-quality checks, allowing researchers to make informed, context-dependent decisions during data preparation.

## REFERENCES

Aiello-Lammens, M. E., Boria, R. A., Radosavljevic, A., Vilela, B., & Anderson, R. P. (2015). spThin: An R package for spatial thinning of species occurrence records for use in ecological niche models. Ecography, 38(5), 541–545. 10.1111/ecog.01132

Boria, R. A., Olson, L. E., Goodman, S. M., & Anderson, R. P. (2014). Spatial filtering to reduce sampling bias can improve the performance of ecological niche models. Ecological Modelling, 275, 73–77. 10.1016/j.ecolmodel.2013.12.012

Castellanos, A. A., Huntley, J. W., Voelker, G., & Lawing, A. M. (2019). Environmental filtering improves ecological niche models across multiple scales. Methods in Ecology and Evolution, 10(4), 481–492. 10.1111/2041-210X.13142

de Lima, R. A. F., Sánchez-Tapia, A., Mortara, S. R., ter Steege, H., & de Siqueira, M. F. (2021). plantR: An R package and workflow for managing species records from biological collections. Methods in Ecology and Evolution, 2021(November), 1–8. 10.1111/2041-210X.13779

Lamboley, Q., & Fourcade, Y. (2024). No optimal spatial filtering distance for mitigating sampling bias in ecological niche models. Journal of Biogeography, 51(9), 1783–1794. 10.1111/jbi.14854

Peterson, A. T., Asase, A., Canhos, D. A. L., de Souza, S., & Wieczorek, J. (2018). Data leakage and loss in biodiversity informatics. Biodiversity Data Journal, 6. 10.3897/BDJ.6.e26826

Reginato, M., & Michelangeli, F. A. (2020). Bioregions of Eastern Brazil, Based on Vascular Plant Occurrence Data. In Neotropical Diversification: Patterns and Processes (pp. 475–494). 10.1007/978-3-030-31167-4_18

Ribeiro, B. R., Velazco, S. J. E., Guidoni-Martins, K., Tessarolo, G., Jardim, L., Bachman, S. P., & Loyola, R. (2022). bdc: A toolkit for standardizing, integrating and cleaning biodiversity data. Methods in Ecology and Evolution, 13(7), 1421–1428. 10.1111/2041-210X.13868

Sillero, N., Arenas-Castro, S., Enriquez□ Urzelai, U., Vale, C. G., Sousa-Guedes, D., Martínez-Freiría, F., Real, R., & Barbosa, A. M. (2021). Want to model a species niche? A step-by-step guideline on correlative ecological niche modelling. Ecological Modelling, 456(July). 10.1016/j.ecolmodel.2021.109671

Telenius, A. (2011). Biodiversity information goes public: GBIF at your service. Nordic Journal of Botany, 29(3), 378–381. 10.1111/j.1756-1051.2011.01167.x

Trindade, W. C. F., & Marques, M. C. M. (2023). The Invisible Species: Big Data Unveil Coverage Gaps in the Atlantic Forest Hotspot. Diversity and Distributions, 30(12), e13931. 10.1111/ddi.13931

Vancine, M. H., Niebuhr, B., Muylaert, R., Galetti, M., & Ribeiro, M. C. (2025). atlanticr: Easy access to data from the ATLANTIC SERIES of data papers for the Atlantic Forest (Version 0.0.0.9000) [R]. https://github.com/mauriciovancine/atlanticr

Velazco, S. J. E., Svenning, J. C., Ribeiro, B. R., & Laureto, L. M. O. (2021). On opportunities and threats to conserve the phylogenetic diversity of Neotropical palms. Diversity and Distributions, 27(3), 512–523. 10.1111/ddi.13215

Wieczorek, J., Bloom, D., Guralnick, R., Blum, S., Döring, M., Giovanni, R., Robertson, T., & Vieglais, D. (2012). Darwin Core: An Evolving Community-Developed Biodiversity Data Standard. PLOS ONE, 7(1), e29715. 10.1371/journal.pone.0029715

Zizka, A., Carvalho, F. A., Calvente, A., Baez-Lizarazo, M. R., Cabral, A., Ramos Coelho, J. F., Colli-Silva, M., Fantinati, M. R., Fernandes, M. F., Ferreira-Araújo, T., Lambert Moreira, F. G., da Cunha Santos, N. M., Borges Santos, T. A., dos Santos-Costa, R. C., Serrano, F. C., da Silva, A. P. A., de Souza Soares, A., de Souza, P. G. C., Tomaz, E. C., … Antonelli, A. (2020). No one-size-fits-all solution to clean GBIF. PeerJ, 8. 10.7717/peerj.9916

Zizka, A., Silvestro, D., Andermann, T., Azevedo, J., Duarte Ritter, C., Edler, D., Farooq, H., Herdean, A., Ariza, M., Scharn, R., Svantesson, S., Wengström, N., Zizka, V., & Antonelli, A. (2019). CoordinateCleaner: Standardized cleaning of occurrence records from biological collection databases. Methods in Ecology and Evolution, 10(5), 744–751. 10.1111/2041-210X.13152

